# Cell Types or Cell States? An Investigation of Adrenergic and Mesenchymal Cell Phenotypes in Neuroblastoma

**DOI:** 10.1101/2023.12.20.572368

**Authors:** Anuraag Bukkuri, Stina Andersson, Joel S. Brown, Emma U. Hammarlund, Sofie Mohlin

## Abstract

Neuroblastoma is a pediatric cancer that exhibits two cellular phenotypes: adrenergic (ADRN) and mesenchymal (MES). ADRN is differentiated and therapy-sensitive, while MES is less differentiated with elevated therapy resistance. To understand neuroblastoma and its treatment response, it is important to elucidate how these phenotypes impact the eco-evolutionary dynamics of cancer cell populations and whether they represent distinct cell types or dynamic cell states. Here, we show that neuroblastoma cells undergo an ADRN to a MES phenotypic switch under chemotherapy treatment. We use a strong inference approach to generate four hypotheses on how this switch may occur: cell types without resistance, cell types with resistance, cell states without resistance, and cell states with resistance. For each of these hypotheses, we create theoretical models to make qualitative predictions about their resulting eco-evolutionary dynamics. Our results provide a framework to further experimentally determine whether ADRN and MES phenotypes are distinct cell types or dynamic cell states.

## Introduction

Neuroblastoma is a malignancy of the sympathetic nervous system. One of the most common and deadliest pediatric cancers, neuroblastoma shows a range of clinical outcomes, from spontaneous regression to metastatic, therapy-resistant cancer with poor patient outcomes. However, the mechanisms underlying the initiation, progression, and emergence of therapeutic resistance in neuroblastoma are not fully understood. Neuroblastoma cells can develop resistance via mutations in specific pathways (primarily the Ras/MAPK signaling pathway^1^). However, it has been shown that mutation-independent phenotypic plasticity in cell state transitions also allow for adaptation to stressful environments.

In recent years, experimental studies using RNA sequencing and epigenetic profiling have found two cancer cell phenotypes in neuroblastoma with divergent gene expression profiles: adrenergic (ADRN) and mesenchymal (MES).^2,3,4,5,6^ The differentiated ADRN phenotype is more sensitive to therapy than the undifferentiated MES phenotype but comprises a higher portion of the population under baseline conditions.^2^ Although patient tumors most commonly present with an ADRN phenotype, using protein markers, epigenetic analyses, or bulk RNA sequencing, studies have identified mesenchymal cells in patients, a fraction of tumors even presenting with a dominant mesenchymal phenotype.^2,7–10^ Furthermore, evidence suggests that cells can switch between these phenotypes, *i*.*e*., a cell with a MES phenotype can adopt an ADRN phenotype and vice versa.^2,8,11^ This trait is described as plasticity. Such interconversion has been demonstrated for several neuroblastoma cell lines.^2,8,12,13^ This phenotypic plasticity, where the fraction of cells with an adopted MES phenotype quickly expand and constitute the majority of the complete cell population in response to chemotherapy or ALK inhibitor treatment,^2,14^ likely plays a key role in the ability of neuroblastomas to develop resistance to therapy.

However, the field lacks a consensus on the classification of these phenotypes. The ADRN and MES phenotypes are sometimes treated as separate cell types, cell states, or the distinction is evaded altogether by classifying cells as ADRN-or MES-like. In fact, these terms are often used interchangeably within the same study. However, it is critical to make a distinction between cell types and cell states, as this has important implications for the eco-evolutionary dynamics of neuroblastoma cell populations under therapy. Cell types refer to heritably distinct cellular species in which progeny resemble their parents; cell states refer to transient phenotypes that cells can adopt and dynamically shift between in a phenotypically plastic manner. Although some studies attempt to develop methods to delineate this difference, they have mainly taken a gene-centric approach to this problem of distinction, omitting broader implications at the population level.^15,16^

In this paper, we address the question of whether ADRN and MES phenotypes represent different cell types or cell states and whether cells with an ADRN phenotype can evolve resistance to therapy using a strong inference approach.^17^ This approach, originally proposed by John Platt in 1964, consists of generating several competing (falsifiable) hypotheses and devising tests that can distinguish between them. This is to ascertain the hypothesis most in accordance with observed reality. In the words of Rob Phillips, we do this “by turning our thinking into formal mathematical predictions and confronting that math with experiments that have not yet been done,” or also described as “Figure 1 Theory”.^18^ To do this, we run preliminary experiments to show the ADRN to MES phenotypic switch that occurs under therapy. We then create mathematical models of four hypotheses to explain how this switch may happen: cell types without resistance (H1), cell types with resistance (H2), cell states without resistance (H3), and cell states with resistance (H4). We use simulations to qualitatively predict trends in the ecological (population) and evolutionary (drug resistance) dynamics of the ADRN and MES phenotypes under treatment. We find that if ADRN cells evolve resistance, the frequency of ADRN cells will increase with subsequent therapeutic insult. We also find that if the ADRN and MES phenotypes are cell states, cells in the MES state will increase in frequency upon induction of therapy but will remain stable or decline shortly after. Conversely, if they are cell types, MES cells will increase in frequency for the entire duration of therapy.

**Figure 1.**
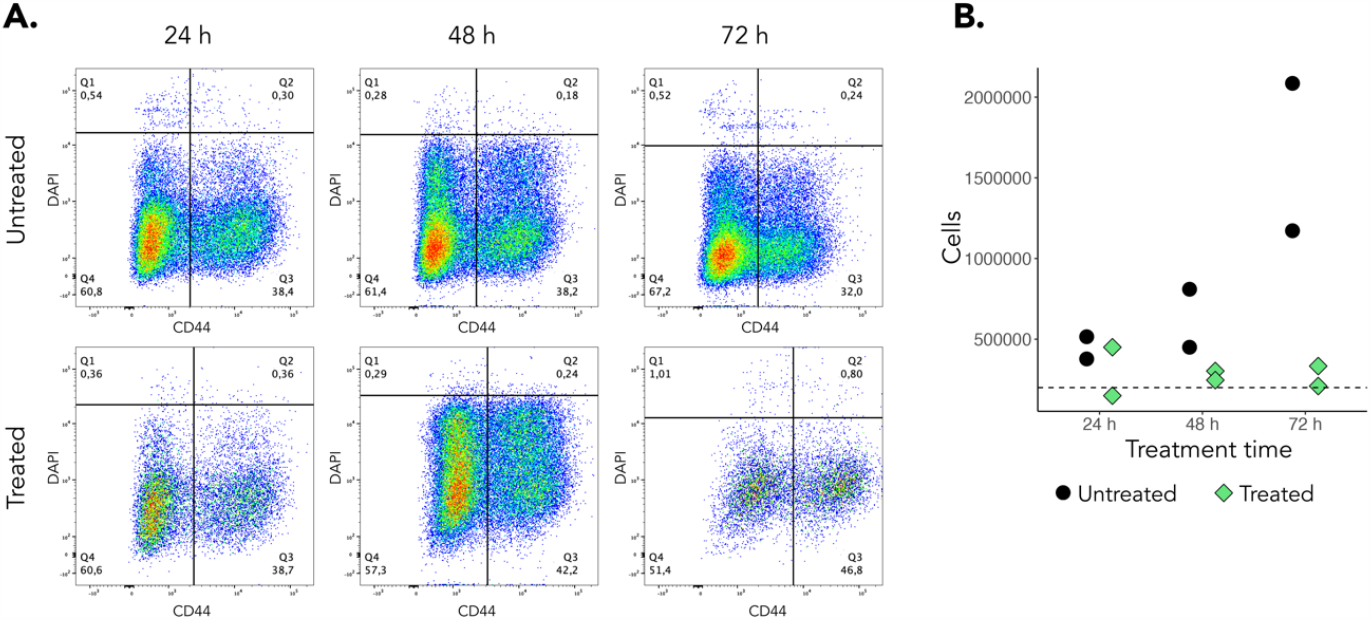
Cisplatin treatment affects CD44 expression and cell growth of SK-N-BE(2) cells. **(A)** Representative dot plots illustrating SK-N-BE(2) cells subjected to treatment for 24, 48 and 72 hours and corresponding untreated cells. Q1 and Q2 represents dead cells, Q3 CD44^high^ cells and Q4 CD44^low^ cells. Three biological repeats were performed. **(B)** Number of viable cells after treatment with 5 μM cisplatin. Dashed line denotes the 200 000 cells seeded 24 hours prior to treatment and diamonds/dots represent two independent biological replicates at each time point.

## Results

### Chemotherapy treatment expands the CD44+ MES cell population

Neuroblastoma cell lines and patient-derived tumors vary in their composition of ADRN- and MES-phenotype cells. Using neuroblastoma SK-N-BE(2) cells, we experimentally determined their ADRN *vs*. MES dynamics at baseline and during chemotherapy treatment. Subclone SK-N-BE(2)c cells have previously been described as primarily presenting with an ADRN phenotype^2^, however, the fraction of each population has not been determined. A recent study identified CD44 as a strong and specific proxy marker for MES cells.^19^ We therefore decided to here use CD44 to sort and distinguish ADRN (CD44^low^) and MES (CD44^high^) cells from each other.

Effects on the ADRN *vs*. MES dynamics after chemotherapy treatment has not been described for this cell line. We treated SK-N-BE(2) cells with cisplatin for 24, 48 or 72 hours, and stained them for CD44 prior to flow cytometry-based analysis. The majority of SK-N-BE(2) cells presented with a dominant ADRN phenotype (CD44^low^) at baseline (Fig. 1A). Our results demonstrate a shift in cell phenotype during treatment, with the fraction of cells expressing mesenchymal CD44 increasing with time exposed to cisplatin (Fig 1A). We also observed that cisplatin-treated cells proliferated at a slower pace (Fig 1B).

Our data are in concordance with previous literature on other neuroblastoma-derived cell lines, and support the notion that neuroblastomas consist of cells of various phenotypes. However, the experimental data do not reveal whether the two phenotypes are distinct cell types or dynamic cell states. To elucidate this question, we turned to mathematical modeling.

### Modeling Framework of Neuroblastoma Eco-Evolutionary Dynamics

To model the eco-evolutionary dynamics of neuroblastoma cancer cell populations, we used an evolutionary game theoretic approach called the G function framework.^20,21^ This framework was originally developed in the context of evolutionary ecology^22–24^ and has recently been applied to problems in cancer.^25–30^ The *G* function, or fitness generating function, is at the core of this framework and captures the per capita growth rate of a population as a function of its mean strategy (*v*) and its density (*x*). With this, we can write the population dynamics of a cancer cell morph as the product of the per capita growth rate and the current population size:

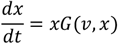

Next, we turned our attention to the strategy dynamics by capturing the resistance level of the morph to a drug. By Fisher’s fundamental theorem,^31–34^ the rate of evolution is proportional to the product of the trait’s evolvability and the strength of selection. This can mathematically be formalized as:

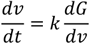

where k is the trait’s evolvability and *dG*/*dv* is the selection gradient, representing how a perturbation in trait value impacts fitness. With this framework, we constructed and simulated *G* function models of the dynamics of neuroblastoma cancer cell populations under therapy. To do this, we considered four hypotheses on the relation between ADRN and MES phenotypes and the evolution of resistance. We created a mathematical model of the underlying ecological and evolutionary dynamics (if applicable). Next, we ran simulations that track the population and resistance strategy dynamics over time, when the population is exposed to continuous and intermittent therapy regimens. The parameter values used in the simulations can be found in Table 1 below and were chosen to parallel qualitative trends observed in the preliminary experiments.

**Table 1.**
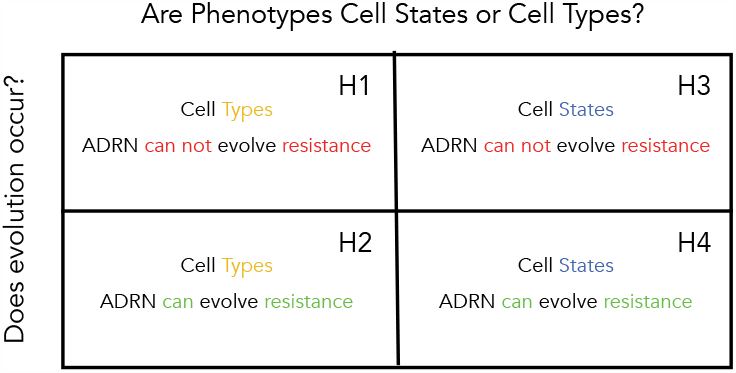
Summary table capturing the four biological hypotheses, surrounding questions of cell type vs. state and evolution of resistance, that will be modeled.

**Table 2:** Parameter Definitions and Values used in Simulations.

| Parameter | Interpretation | Value |
| --- | --- | --- |
| $r_A$ | Intrinsic Growth Rate | 0.6 |
| $r_M$ | Intrinsic Growth Rate of MES Cells | 0.6 |
| $K$ | Carrying Capacity | $5 * 10^6$ |
| $\gamma$ | Spontaneous Transition Rate from ADRN to MES | 0.02 |
| $m$ | Drug Dosage | 0.1 |
| $\lambda$ | Baseline Level of Resistance | 1 |
| $b$ | Efficacy of Evolving a Resistance Strategy | 1 |
| $a$ | Facultative Transition Scaling Rate from ADRN to MES | 0.8 |
| $\mu$ | Spontaneous MES to ADRN Transition Rate | 0.2 |
| $\alpha$ | Competitive Effect of MES cells on ADRN Cells | 0.95 |
| $\beta$ | Competitive Effect of ADRN cells on MES Cells | 0.98 |
| $k$ | Evolvability | 0.4 |
| $v$ | Drug Resistance | $[0, \infty)$ |

**Table 3.** List of antibodies.

| Antibody | Species | Dilution | Source | Product # |
| --- | --- | --- | --- | --- |
| Human/Mouse CD44 Alexa Fluor® 647-conjugated Antibody | Rat | 1:50 | R&D Systems | FAB6127R |
| Rat IgG2B Alexa Fluor® 647-conjugated Isotype Control | Rat | 1:50 | R&D Systems | IC013R |

### Cell Type without Resistance Hypothesis

The first hypothesis (H1, Table 1) views the ADRN and MES phenotypes as distinct cell types and assumes that ADRN cells remain sensitive to therapy without evolving resistance. Since ADRN and MES cells in this scenario are different cell types, we do not allow interconversion between them. We assumed that the cells grow in a logistic manner. Since it is not experimentally feasible to isolate ADRN and MES cells and measure intrinsic growth rates, we assume ADRN and MES cells have the same proliferation rate, but imbue ADRN cells with a competitive advantage over MES cells (α < β) to ensure that they attain a higher frequency in the population at baseline. Although there are alternative ways of doing this, such as by

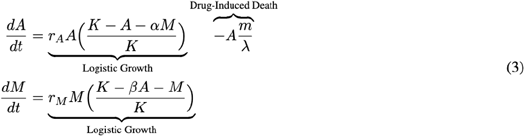

allowing ADRN cells to have the same intrinsic death rate and a higher intrinsic growth rate than MES cells, they do not alter qualitative results. For simplicity, we assumed that MES cells are fully resistant to therapy and ADRN cells die in a constant, density independent manner from therapy. Since we assume ADRN cells do not evolve resistance, we do not have any evolutionary dynamics in this model. Mathematically, we can formalize this hypothesis as follows:

With this model, we further simulated therapy, which we chose to administer from time 300 to 400 and time 1200 to 1300 (Fig. 2). Drug holidays allow the population to recover to near-homeostatic conditions. Under both regimens, ADRN cells have a competitive fitness advantage over MES cells under no therapy, and thus comprised most of the population under baseline conditions. Indeed, the 71% ADRN 29% MES frequency under untreated homeostasis (as determined by comparing the proportions of ADRN and MES cells in our model simulation (Fig. 2) immediately before therapeutic periods) is in accordance with experimental observations (Fig. 1A). During times of therapy, MES cells expand and outcompete ADRN cells since MES cells are agnostic to therapy and ADRN cells are sensitive. Furthermore, since ADRN cells do not gain resistance to therapy, repeated therapeutic insults gave identical ecological responses, and the same cyclical behavior was observed across on-off cycles.

**Figure 2.**
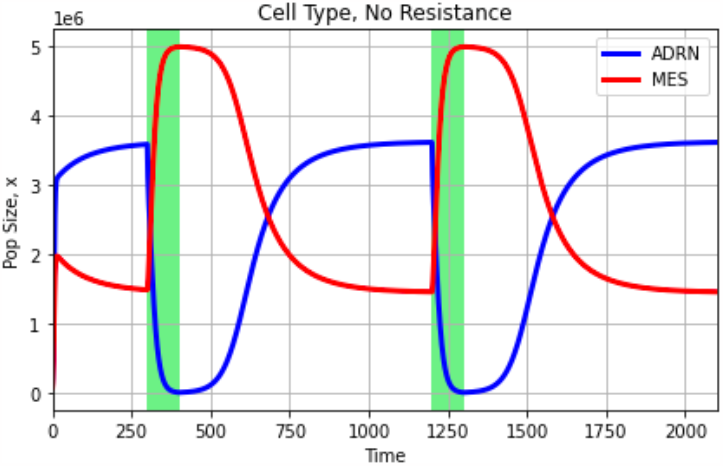
Simulations under Cell Type with Resistance Hypothesis. Blue and red curves capture ADRN and MES population dynamics. White and green backgrounds reflect periods without and periods with treatment. During periods of therapy, MES cells have a competitive advantage over ADRN cells, leading to an increase in number and frequency. When therapy is removed, the opposite is true, with ADRN cells taking over the population. Since ADRN cells do not gain resistance, identical ADRN and MES population dynamics are observed for repeated bouts of therapy.

It’s worth noting that, particularly under therapy, the parameter values we choose can influence which cell type dominates the population. For instance, it could be expected that higher drug dosages would promote the expansion of MES cells in the population. To investigate this idea, we analytically compute the equilibria of our model under therapy:

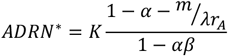

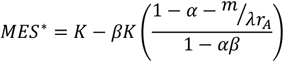

Notice the impact of drug dosage and ADRN proliferation rate on the ADRN and MES equilibria: drug dosage decreased (increased) the ADRN (MES) equilibrium, whereas ADRN proliferation rate increased (decreased) the ADRN (MES) equilibrium. This can be visualized by plotting the equilibria as a function of drug dosage and ADRN proliferation rate (Fig. 3).

**Figure 3.**
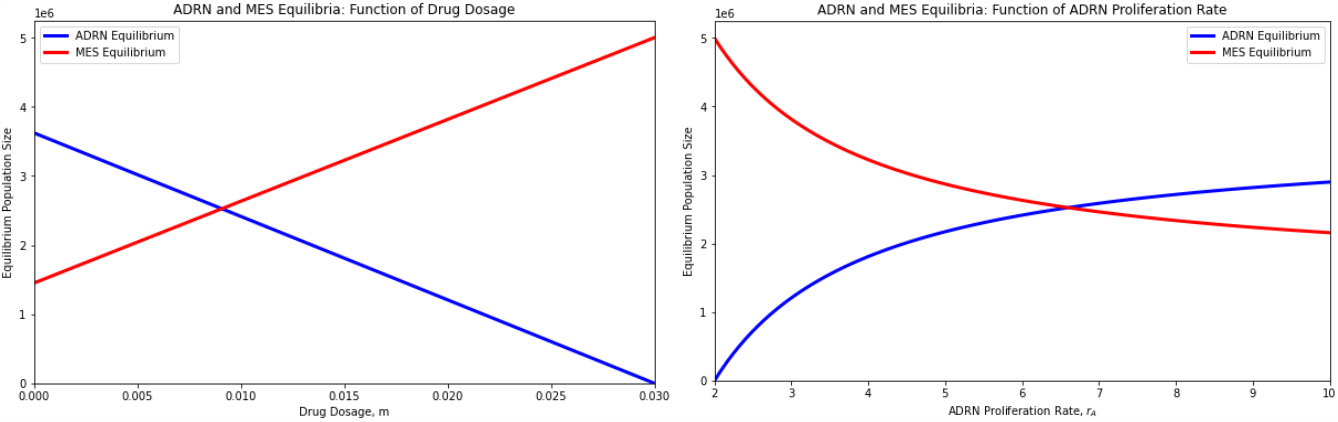
ADRN and MES Equilibria Plotted as Functions of Drug Dosage and ADRN Proliferation Rate. Blue and red curves capture ADRN and MES equilibria. The left (right) plot is produced by varying *m*(*r*_*A*_) and setting all other parameters to their default values, as given in Table 1. Drug dosage decreases the ADRN equilibrium and increases the MES equilibrium. Conversely, ADRN proliferation rate increases the ADRN equilibrium and decreases the MES equilibrium.

### Cell Type with Resistance Hypothesis

For our second hypothesis (H2, Table 1), we still assumed that the ADRN and MES phenotypes represent different cell types, but we allowed ADRN cells to evolve resistance to therapy. To do this, we modified

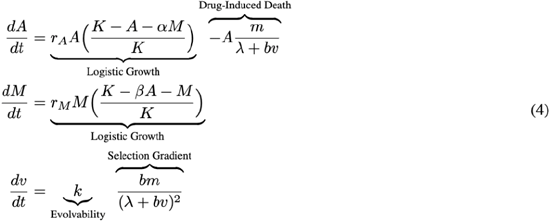

our drug-induced death term to account for the impact of a resistance strategy and added an equation for the evolutionary dynamics of drug resistance. Note that we assumed that ADRN cells suffer no cost from evolving resistance to Cisplatin. Since ADRN and MES cells in this scenario are still different cell types, resistance is a trait of solely ADRN cells. Thus, the evolutionary dynamics are derived only from the ecological dynamics of ADRN cells as shown under the ‘Modeling Framework’ section.

We simulated therapy, administered from time 300 to 400 and 1000 to 1100, where drug holidays allow cells to return to near homeostatic conditions (Fig. 4). Overall, we noticed similar dynamics to our previous hypothesis: ADRN cells outcompete MES cells when no therapy is present and MES cells have an advantage during times of therapy. As expected, periods of therapy promote the evolution of the strategy toward higher resistance levels, while periods without therapy induce evolutionary stasis. Note that the rate of evolution of resistance of ADRN cells decreases as cells acquire resistance. This is because their selection gradient (selection pressure) decreases, thereby slowing the evolution of resistance (see Equation 2). Furthermore, since ADRN cells gain resistance to therapy over time, the MES cells have less of an advantage during each repeated therapeutic insult. This leads to a smaller increase (decrease) in MES (ADRN) cells during each therapeutic period.

**Figure 4.**
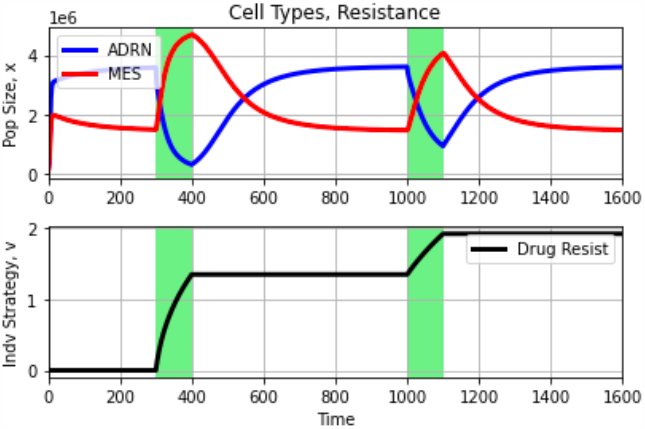
Simulations under Cell Type with Resistance Hypothesis. Blue and red curves capture ADRN and MES population dynamics. The black curve represents the evolutionary dynamics of resistance. White and green backgrounds reflect periods without and periods with treatment, respectively. During periods of therapy, MES cells have a competitive advantage over ADRN cells, leading to an increase in number and frequency. When therapy is removed, the opposite is true, with ADRN cells taking over the population. As ADRN cells gain resistance, the relative advantage of MES cells during periods of therapy decreases.

Since we did not include a cost of resistance, there is ostensibly a level of resistance for which MES cells no longer outcompete ADRN cells in the presence of therapy. To find this point, we analytically solve for the ecological equilibria of our system:

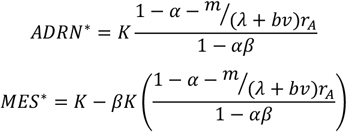

We again plotted the ADRN and MES equilibrium values as a function of drug resistance, *v* (Fig. 5). For our parameter values, the ADRN ecological equilibrium surpasses the MES equilibrium when they attain a resistance value of v>10. In other words, in a treated environment in which m=0.1, if v>10, the population will primarily be composed of ADRN cells at equilibrium, whereas if v<10, the population will primarily be composed of MES cells at equilibrium.

**Figure 5.**
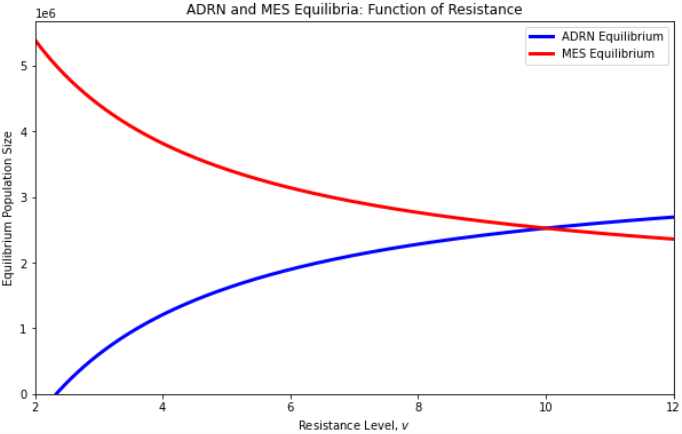
ADRN and MES Equilibria Plotted as Functions of Resistance Level. Blue and red curves capture ADRN and MES equilibria. The plot is produced by varying 𝒱 and setting all other parameters to their default values, as given in Table 1. Higher drug resistance levels promote higher ADRN equilibria and lower MES equilibria.

### Cell State without Resistance Hypothesis

Our third hypothesis (H3, Table 1) considers the ADRN and MES phenotypes as different cell states in the life cycle of a cell (species) and assumes that ADRN cells do not evolve resistance to therapy. This means that we allow for interconversion between cells in the ADRN and MES states. Specifically, we allow for a

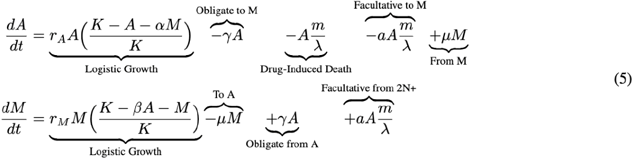

spontaneous, obligate, background transition between cell states as well as a facultative, condition-dependent transition from the ADRN to MES state. When cells are under stress due to therapy, they can transition to the MES state, which serves as a refuge from therapy. This can be mathematically formalized as shown in Equation 5. Note that since we do not allow for the evolution of resistance, we are only concerned with ecological dynamics in this situation.

Next, we simulated therapy, administered from time 100 to 200 and 300 to 400 (Fig. 6). During periods of no therapy, we noticed that cells reach and maintain an equilibrium composed of primarily cells in the ADRN state. As before, the 71% ADRN and 29% MES frequency under baseline conditions parallel experimental results (Fig. 1A). Under therapy, cells rapidly switch to the MES state to avoid the effects of therapy. The population of cancer cells then exists primarily in a MES state. This can be conceptualized as a transient state of partial resistance in the population since cells do not actually acquire resistance but instead transition in a plastic manner to a MES state of resistance, and not all cells exist in the MES state during therapy. Because we do not allow for the evolution of resistance, repeated therapeutic administrations lead to identical ecological dynamics.

**Figure 6.**
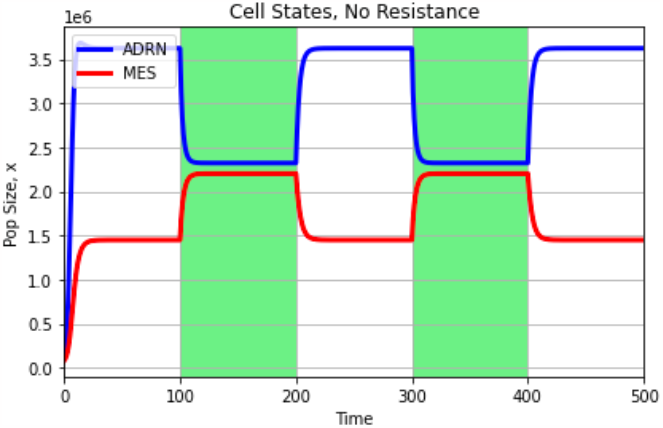
Simulations under Cell State without Resistance Hypothesis. Blue and red curves capture ADRN and MES population dynamics, respectively. White and green backgrounds reflect periods without and periods with treatment. Upon administration of therapy, we see a dramatic shift in frequency and number from the ADRN state to the MES state, allowing the population to seek refuge from therapy. However, since cells in the ADRN state do not evolve resistance, this represents a temporary state of resistance. After therapy is removed, the population reverts to a primarily sensitive phenotype, with most cells transitioning back to the ADRN state.

### Cell State with Resistance Hypothesis

Our fourth hypothesis (H4, Table 1) views ADRN and MES phenotypes as cellular states and allows for the evolution of resistance. Our ecological dynamics are similar to before, although we modify our drug-induced death and facultative transition terms to account for the impact of resistance. Since the ADRN and MES phenotypes are conceptualized as states in the life history of the same cell type instead of as separate cell types altogether, a unifying fitness function across both cell states is required. To do this, we utilize a method developed^35^ and applied^26,29^ in prior work that combines techniques from evolutionary game theory and matrix population modeling. We represent our ordinary differential equation model as a population projection matrix (PPM) and use the spectral bound (ρ(PPM)) of this matrix as a measure of fitness, as it controls the long-term, asymptotic growth rate of the population. By setting G = ρ(PPM), we derived our evolutionary dynamics as outlined under the ‘Modeling Framework’ section.

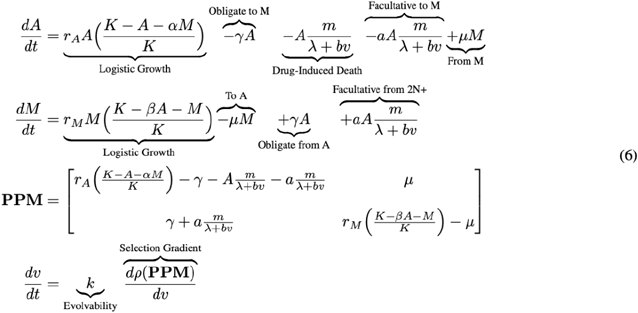

With this model, we simulated the effects of therapy, administered from time 100 to 200 and time 300 to 400 (Fig. 7). As before, during times of no therapy, the population rapidly reaches and maintains an equilibrium of cells primarily in the ADRN state. When therapy is administered, the number and frequency of cells in the MES state increase dramatically. However, as cells become increasingly resistant, the death due to drug and facultative transitions to the MES state decreases as well. Consequently, we note a gradual transition of cells from the MES state to the ADRN state during therapy. Note that the dramatic shift of cells to the MES state followed by a gradual transition of cells to the ADRN state under therapy is unique to this hypothesis.

**Figure 7.**
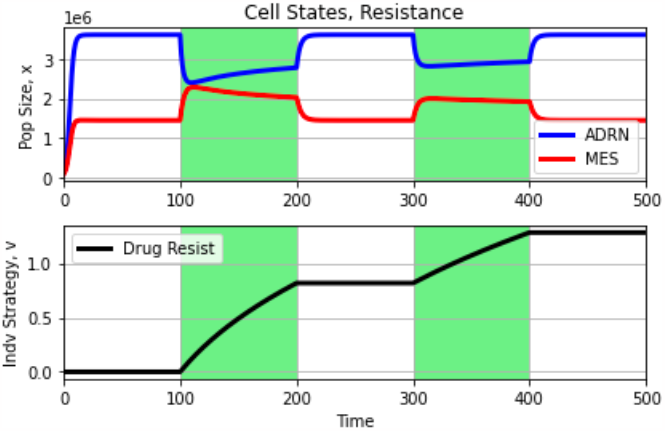
Simulations under Cell State with Resistance Hypothesis. Blue and red curves capture ADRN and MES population dynamics, respectively. The black curve represents the evolutionary dynamics of resistance. White and green backgrounds reflect periods without and periods with treatment. Upon administration of therapy, we see a dramatic shift in frequency and number from the ADRN state to the MES state, allowing the population to seek refuge from therapy. As cells in the ADRN state gain resistance, the frequency and number of cells in the ADRN (MES) state increases (decreases). When therapy is removed altogether, the population reverts to its pre-therapy distribution, with most cells existing in the ADRN state.

## Discussion

The finding in 2017^2^ that neuroblastomas are composed of two distinct phenotypes, ADRN and MES, has shaped how the field views clinical outcomes and has influenced pre-clinical experimental design and analysis. While many chemotherapeutics and targeted therapies used for neuroblastoma patients are successful in the initial phase, the treatment often fails eventually. In order to overcome this issue, it is critical to understand the ecological and evolutionary dynamics of cellular populations with ADRN and MES phenotypes, particularly under therapy. In line with previously published data, we experimentally show here that neuroblastoma cell populations treated with chemotherapy *in vitro* shift from an ADRN phenotype-dominated composition to a MES phenotype. However, whether such a shift is driven by demographic transitions between cell states or is the result of ecological interactions between cell types is unknown. Furthermore, whether cells with the ADRN phenotype can evolve resistance is unclear.

We used our generated cell population data to test four different hypotheses addressing the concepts of cell type versus cell state and evolution of resistance of the ADRN phenotype. We created eco-evolutionary mathematical models for each hypothesis and qualitatively simulated expected ecological, evolutionary, and demographic dynamics under therapy. Our simulations showed that if cells with an ADRN phenotype can evolve resistance to therapy, we would notice different dynamics with repeated administrations of therapy.

Namely, the frequency of cells with a ADRN phenotype would increase with subsequent therapeutic insults; while the MES phenotype would decrease. Conversely, for resistance to not evolve requires that population dynamics are identical across therapeutic cycles. Also, our simulations show that if the ADRN and MES phenotypes represent cell types, cells of a MES phenotype would increase in frequency for the entire duration of the therapeutic period. If they are instead cell states, cells of a MES phenotype would increase in frequency initially but will either begin a gradual decline soon after (if evolution of resistance occurs) or will remain stable (if evolution of resistance does not occur) during periods of therapy. In summary, our mathematical models, inspired by preliminary biological data, provide a framework for further experimental steps to elucidate the characteristics of neuroblastoma cell phenotype (state or type), the evolution of resistance, and how these impact treatment response.

### Limitations of the study

The experimental part of this study is limited to *in vitro* experiments in one cell line. Although our experiments corroborate with previously published data, our results should be validated in additional cell lines and in an *in vivo* setting. Additionally, our model assumptions, notably the lack of a cost of resistance, should be validated with *in vitro* experiments.

## Acknowledgements

This study was funded by Längmanska kulturfonden, the Royal Swedish Academy of Sciences, GS Magnuson Foundation, the National Science Foundation Graduate Research Fellowship Program, the Swedish Cancer Society, The Swedish Childhood Cancer Fund, The Crafoord Foundation, The Ollie and Elof Ericsson Foundation, The Magnus Bergvall Foundation, The Hans von Kantzow Foundation, The Royal Physiographic Society of Lund. The authors would like to thank Anna Hammarberg and the FACS Core Facilities at Multipark and Lund Stem Cell Center for technical expertise.

## Author contributions

AB and SM conceptualized the article. AB and JSB developed the models and ran the simulations. AB, SA, and SM designed the experiments. SA performed experiments and analyzed the experimental data. AB, SA, and SM wrote the original draft of the paper. All authors edited and approved the final version of this article.

## Data Availability

All data generated or analyzed during this study are included in this published article.

## Material and methods

### Cell culture

The neuroblastoma cell line SK-N-BE(2) (ATCC, 2022) was cultured in MEM supplemented with 10% fetal bovine serum, 100 units penicillin and 10 μg/mL streptomycin. Cells were kept at 37°C, 21% O_2_ and 5% CO_2_ in a humidified incubator and dissociated with trypsin. Cells were routinely tested for mycoplasma.

### Cell counting

Cells were stained with Trypan blue and live cells were counted in two technical replicates with a TC20 Automated Cell Counter (Bio-Rad).

### Flow cytometry

200 000 cells were seeded to 35 mm wells and were either treated with 5 μM cisplatin after 24 hours or kept as untreated control. Cells were harvested with trypsin after 24, 48 or 72 hours of treatment, washed once in PBS and stained with antibodies (Table 1) in 100 μL FACS buffer (PBS, 0.5% BSA, 4mM EDTA) at 4°C avoiding light. Cells were then washed once again in PBS, resuspended in FACS buffer with 1:3000 DAPI and subjected to flow cytometry in BD LSRII or BD LSR Fortessa. Compensation controls, FMO controls and isotype controls were included in each run and samples were run in three technical replicates.

### Data analysis

Flow cytometry data was analyzed with FlowJo v10. Cell debris and duplicate cells were excluded by gating SSC-A against FSC-A followed by FSC-A against FSC-W. Gating had to be adapted to each biological replicate as the autofluorescence varied between them.

